# Dynamic CRMP2 regulation of CaV2.2 in the prefrontal cortex contributes to the reinstatement of cocaine seeking

**DOI:** 10.1101/533083

**Authors:** William C. Buchta, Aubin Moutal, Bethany Hines, Constanza Garcia-Keller, Alexander C.W. Smith, Peter Kalivas, Rajesh Khanna, Arthur C. Riegel

**Affiliations:** Department of Neuroscience, Medical University of South Carolina, Charleston, South Carolina, 29425; Neurobiology of Addiction Research Center, Medical University of South Carolina, Charleston, South Carolina, 29425.; Department of Pharmacology, University of Arizona, Tucson, Arizona, 85724; Department of Anesthesiology, University of Arizona, Tucson, Arizona, 85724; The Center for Innovation in Brain Sciences, The University of Arizona Health Sciences, Tucson, Arizona

**Keywords:** CaV2.2, cocaine reinstatement, collapsin response mediator protein 2 (CRMP2), myr-TAT-CBD3, prefrontal cortex (PFC)

## Abstract

Cocaine addiction is a major health concern with limited effective treatment options. A better understanding of mechanisms underlying relapse may help inform the development of new pharmacotherapies. Emerging evidence suggests that collapsin response mediator protein 2 (CRMP2) regulates presynaptic excitatory neurotransmission and contributes to pathological changes during diseases, such as neuropathic pain and substance use disorders. We examined the role of CRMP2 and its interactions with a known binding partner, CaV2.2, in cocaine-seeking behavior. We employed the rodent self-administration model of relapse to drug-seeking and focused on the prefrontal cortex (PFC) for its well-established role in reinstatement behaviors. Our results indicated that repeated cocaine self-administration resulted in a dynamic and persistent alteration in the PFC expression of CRMP2 and its binding partner, the CaV2.2 (N-type) voltage-gated calcium channel. Following cocaine self-administration and extinction training, the expression of both CRMP2 and CaV2.2 was reduced relative to Yoked saline controls. By contrast, cued-reinstatement potentiated CRMP2 expression and increased CaV2.2 expression above extinction levels. Lastly, we utilized the recently developed peptide *myr-TAT-CBD3* to disrupt the interaction between CRMP2 and CaV2.2 *in vivo*. We assessed the reinstatement behavior after infusing this peptide directly into the medial PFC and found that it decreased cue-induced reinstatement of cocaine seeking. Taken together, these data suggest that neuroadaptations in the CRMP2/CaV2.2 signaling cascade in the PFC can facilitate drug seeking behavior. Targeting such interactions has implications for the treatment of cocaine relapse behavior.

## Introduction

Cocaine addiction is a major health concern with limited effective treatment options. Currently, there are no effective medications approved to treat cocaine addiction. Despite the recognition of addiction as a disease, high relapse rates remain a major challenge for the treatment of cocaine addiction and other substance abuse disorders. A better understanding of the mechanisms underlying relapse may help inform the development of improved pharmacotherapies.

In humans and preclinical rodent models, relapse can occur long after the cessation of drug taking and is often precipitated by exposure to drug-associated cues [15]. A prevailing hypothesis is that this behavior reflects a drug-induced maladaptive rewiring of cortical-striatal circuitry in the medial prefrontal cortex (mPFC) and nucleus accumbens regions [19; 20]. The persistence of this plasticity and related behaviors may involve neuroadaptations in synaptic strength [9] and gene expression [14; 40], as well as cytoskeletal re-arrangement [13]. Conversely, disrupting these adaptations reduces drug-induced alterations in excitability and associated behaviors [38]. It is unclear if these abnormalities are a precedent or an antecedent of addiction.

Potential substrates in addiction etiology may include enduring adaptations in presynaptic glutamate release in the PFC [31; 46]. Addictive drugs can both sensitize glutamate release in the mPFC [31; 46; 47] and upregulate the auxiliary α2δ1 subunit of CaV2.2 (N-type) and CaV2.1 (P/Q-type) voltage-gated calcium channels (VGCCs) [21; 28; 45]. Another protein regulating CaV2.2, the axonal growth/guidance collapsin response mediator protein 2 (CRMP2) [8; 22], was found to be associated with the molecular pathology underlying addictive behaviors [6; 29]. CRMP2 is an intracellular protein participating in growth cone collapse induced by semaphorin3A and axonal growth by promoting tubulin polymerization [53]. CRMP2 regulates the trafficking and the presynaptic localization of the voltage gated calcium channel CaV2.2, an association that is essential for excitatory spinal glutamatergic neurotransmission [10; 33; 35; 36].

After binge alcohol drinking, CRMP2 mRNA translation was increased which lead to increased protein expression level at the synaptic sites in the nucleus accumbens [4; 30]. This newly synthesized CRMP2 existed in a form not phosphorylated by glycogen synthase kinase 3β (GSK-3β) and showed elevated binding to microtubules [4; 30]. Chronic cocaine or methamphetamine exposure can decrease CRMP2 abundance [25; 50] and long-term withdrawal from cocaine can increase CRMP2 phosphorylation level by protein kinase A [6]. Moreover, a priming injection of alcohol sufficient to reinstate conditioned place preference increased striatal CRMP2 abundance, and conversely systemic administration of a CRMP2 inhibitor decreased excessive alcohol intake [4; 30].

The role of CRMP2 in PFC function and drug seeking remains less clear. At 1 day after acute methamphetamine exposure, CRMP2 abundance in the cortex was decreased [27]. However, to the best of our knowledge, no studies have examined CRMP2 function in the PFC circuitry in the context of drug seeking. Based on findings in the literature, we tested whether chronic cocaine self-administration and extinction would decrease CRMP2 abundance in the PFC and to what extent drug-paired cues (without drug delivery) would alter CRMP2-CaV2.2 expression in the PFC. Lastly, we tested the prediction that interfering with this interaction, with a peptide specifically blocking CRMP2/CaV2.2 interaction, would prevent cocaine seeking behaviors. Our results indicate that cocaine self-administration and reinstatement dynamically modify PFC expression of CRMP2 and its binding partner CaV2.2, and that these complex neuroadaptations may be necessary for cocaine seeking behavior.

## Materials and Methods

All methods used followed National Institutes of Health guidelines for care of laboratory animals and were approved by the Medical University of South Carolina Internal Animal Care and Use Committee. Male Sprague-Dawley rats were single housed with a 12-hour reverse light/dark cycle and were acclimated to the vivarium with access to food and water ad libitum for at least a week prior to surgery.

### Peptide

The N-terminal myristoylated peptide (myr-tat-CBD3; N-myristoyl-<u>YGRKKRRQRRR</u>ARSRLAELRGVPRGL; tat sequence denoted in the underlined text), was synthesized and HPLC-purified by Genscript USA Inc. (Piscataway, NJ). This peptide was characterized extensively in previous studies [18] and found to inhibit specifically the interaction between CRMP2 and CaV2.2. Previously, we also showed that the TAT cell penetrating motif (CPM), the CBD3 peptide without the TAT-CPM (without the ability to enter the cell) and two myristoylated TAT-fused scrambled controls did not show any activity on CaV2.2 currents or behavior, thus demonstrating that the observed effects of the peptide are entirely due to on-target activity [18].

### Surgery

Surgical procedures included implanting catheters for intravenous cocaine infusions and intra-cranial canulae for injection of pharmacological compounds directly to the mPFC, as previously described [38]. Rats were anaesthetized using a ketamine HCl / xylazine mixture (0.57 / 0.87 mg/kg, respectively, i.p.) followed by ketorolac (2.0 mg/kg, i.p.) and Cefazolin (40 mg, i.p. or 10 mg / 0.1 ml, i.v.). Subsequently, intra-jugular catheters were implanted as described elsewhere [49]. Some rats also received bilateral canulae (double 28 ga barrel; 1.2-1.5 mm C-C; Plastics One) stereotactically implanted into the brain 1 mm above the prelimbic PFC using the following coordinates relative to Bregma (in mm; AP: +3.0-3.5; DV: −3.0; ML: ±0.75).

### Behavioral training and reinstatement testing

Following recovery from surgery, rats were trained to self-administer cocaine for 2 hours daily for 14 days. Rats self-administered cocaine on a fixed ratio 1 (FR1) schedule with a 20-second timeout between infusions. Active lever presses resulted in an 0.2 mg/kg/infusion of cocaine and a 5-second light/tone cue. Following 14 days of meeting self-administration criteria (≥10 infusions), rats underwent 7-14 days of extinction where lever presses had no programmed consequences.

After at least 7 days of extinction training and ≤25 active lever presses in the last two days, some rats were sacrificed for tissue collection and biochemical analysis, while others underwent reinstatement testing prior to sacrifice for tissue collection and biochemical analysis. For these rats, reinstatement testing consisted of a 30 min cue-reinstatement session, where active lever pressing results in a 5-second light/tone cue, but no cocaine infusion. These rats were then immediately sacrificed and mPFC tissue was harvested for biochemical analysis.

A second group of cannulated rats trained and extinguished of cocaine seeking as described above were destined for mPFC drug infusions prior to cocaine-(10 mg/kg, i.p.) or a cue-induced reinstatement testing sessions. These reinstatement sessions lasted 2 hours. During the cocaine-induced reinstatements, lever pressing had no programmed consequences. During the cued reinstatements, active lever pressing results in a 5-second light/tone cue, but no cocaine infusion. Each rat in this group underwent two reinstatement test sessions for both types of reinstatement, counterbalanced with respect to infusion of either myr-TAT-CBD3 (30 μM) or vehicle (0.1% dimethyl-sulfoxide [DMSO] in 0.5 μl saline) into the mPFC 10 mins prior to the start of the 2-hour session.

### Locomotor activity

At least 24 hours after the fourth reinstatement test, rats underwent locomotor testing in a novel chamber following intracranial infusion of either myr-TAT-CBD3 (30 μM) or vehicle (0.1% dimethyl-sulfoxide [DMSO] in 0.5 μl saline). For this, rats received microinjections immediately before placement in the photocell chamber apparatus, to record exploratory movement for 1 hr using software (AccuScan Instruments) that estimated distance traveled based on consecutive breaking of adjacent photobeams. Afterwards, animals were sacrificed for histological verification of canulae placement.

### Generation of lysates

mPFC lysates (including the anterior cingulate, prelimbic area, and infralimbic are) prepared from indicated rats were generated by homogenization and sonication in RIPA buffer (50 mM Tris-HCl, pH 7.4, 50 mM NaCl, 2 mM MgCl_2_, 1% (vol/vol) NP40, 0.5% (mass/vol) sodium deoxycholate, 0.1% (mass/vol) SDS) as previously described [10]. The RIPA buffer included freshly added protease inhibitors (Cat# B14002, Biotools, Houston, TX), phosphatase inhibitors (Cat# B15002, Biotools, Houston, TX), and Benzonase (Cat# 71206, Millipore, Billerica, MA). Protein concentrations were determined using the BCA protein assay (Cat# PI23225, Thermo scientific).

### GST-pull-down and Western blotting

Glutathione magnetic beads (Cat# B23702, Biotools, Houston, TX), pre-incubated with bacteria lysate were CRMP2-GST expression was induced and incubated overnight with 300 μg of total protein from mPFC lysates at 4°C in the absence or presence of the indicated peptides with gentle rotation. Beads were washed 3 times with RIPA buffer before re-suspension in Laemmli buffer and denaturation (5 min at 95°C) and immunoblotting as described previously [7; 26; 52] using validated antibodies. Immunoblots were revealed by enhanced luminescence (WBKLS0500, Millipore) before exposure to photographic film. Films were scanned, digitized, and quantified using Un-Scan-It gel version 6.1 scanning software (Silk Scientific Inc, Orem, UT).

### Statistical analyses

All values represent the mean ± S.E.M., unless noted otherwise. All data was first tested for a Gaussian distribution using a D’Agostino-Pearson test (Graphpad Prism 7 Software). The statistical significance of differences between means was determined by either parametric or non-parametric Student’s t test or ANOVA using GraphPad Prism 7 Software. Differences were considered significant if p≤ 0.05. All data were plotted in GraphPad Prism 7. No outlier data was removed.

## Results

We employed the self-administration model of relapse to drug-seeking [5] to examine the role of PFC CRMP2 and its interaction with CaV2.2 in cocaine-seeking behavior. See Figure 1 for experimental design and timeline. Rats were trained to self-administer cocaine as previously described by us and others [38; 43; 49]. During the 14 days of self-administration training, presses on the active lever resulted in cocaine infusions (0.20 mg/kg/infusion), paired with light/tone cues (**Fig. 2A**). Parallel control rats received yoked infusions of saline (**Fig. 2B**). Following self-administration training, all rats underwent extinction training for 7 to 14 days where neither cues nor drug was available (**Fig. 2A,B**). A subset of cocaine-experienced (n=7) and saline experienced (n=4) rats were sacrificed after extinction for biochemical analysis of mPFC CRMP2 and CaV2.2 proteins. Another subset of rats trained and extinguished of cocaine seeking underwent a 30 min cue-reinstatement session to model relapse to cocaine seeking (n=11), where cocaine-associated cues were presented, but lever pressing did not result in drug delivery (**Fig. 1**). Exposure to cocaine-associated cues potentiated lever pressing in rats with a history of cocaine self-administration and extinction training relative to matched Yoked saline control rats (t_(19)_=6.074, p<.0001) (**Fig. 2A**). Following testing, these rats were then immediately sacrificed for PFC tissue analysis. For controls, PFC tissue was harvested from paired Yoked saline + extinction rats (n=10). The remaining rats trained and extinguished of cocaine-seeking (n=9) were destined for peptide infusion directly into the mPFC prior to reinstatement testing, as shown in Fig. 5.

**FIGURE 1:**
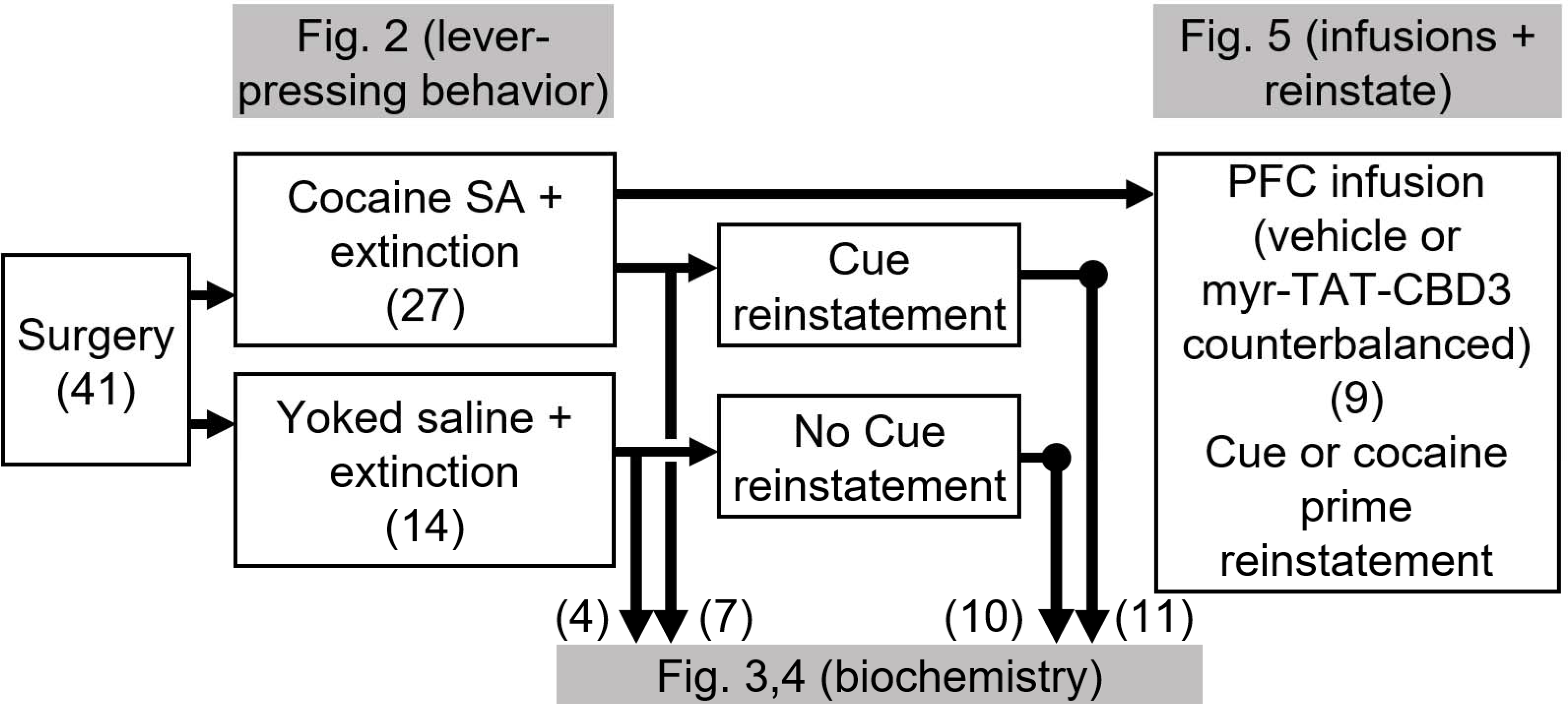
Experimental design and timeline. All rats (n=41) underwent surgery prior to behavioral training. A total of 27 rats underwent cocaine self-administration (SA) training, whereas 14 rats received Yoked saline as a control. All rats then underwent extinction sessions. Subsequently, some rats from each group were sacrificed for biochemical tissue analysis (Cocaine SA + extinction, n=7; Yoked saline, n=4). A second set of cocaine-experienced rats were re-exposed to drug-conditioned cues for 30 mins and then immediately sacrificed for biochemical tissue analysis (Cue reinstatement, n=11) while the remaining Yoked saline rats (n=10) served as controls. The remaining cannulated rats (n=9) trained and extinguished of cocaine seeking were destined for peptide infusion directly into mPFC prior to reinstatement testing, as shown in Fig 5.

**FIGURE 2:**
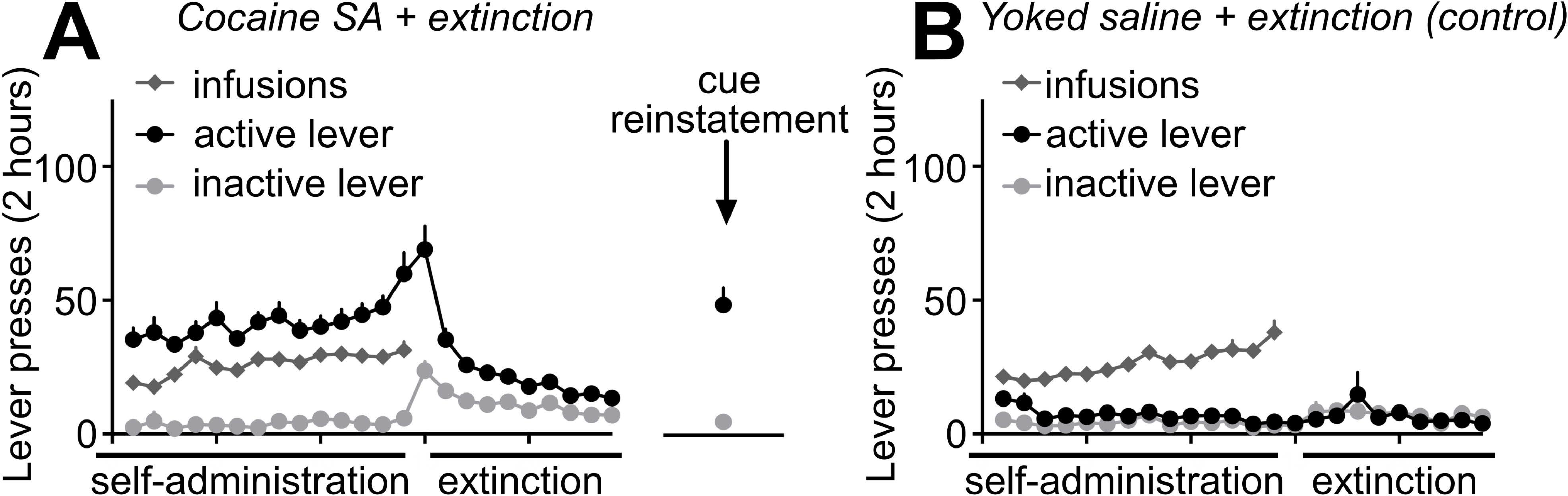
Rodent model of Cocaine SA, extinction training, and reinstatement testing. (**A,B**) All behavioral training for (**A**) Cocaine SA + extinction or (**B**) Yoked saline infusions (control) occurred in operant boxes equipped with inactive levers and active levers. In (**A**), presses on the active lever resulted in an infusion of cocaine paired with the presentation of a light and tone cue. In (**B**), a second group of Yoked control rats received concurrent passive infusions of saline (Yoked saline). After 14 days of Cocaine SA (or Yoked saline), all rats underwent extinction sessions, during which lever pressing produced no infusions or cue presentations. Relative to extinction responding, re-exposure to cocaine-associated cues for 30 mins potentiated lever pressing in rats with a history of Cocaine SA (t_(10)_=5.222, p=0.0004). For these and all other subsequent figures, error bars indicate mean ± SEM.

### CRMP2 expression, but not phosphorylation level, is altered in the mPFC of cocaine self-administration rats

To evaluate whether CRMP2 expression or its phosphorylation level could be altered during reinstatement of cocaine seeking, we performed a Western blot on mPFC tissues from the indicated rats (**Fig. 3**). CRMP2 can be phosphorylated by at least 5 kinases (Fyn, Yes, cyclin-dependent kinase 5 (Cdk5), glycogen synthase kinase 3β (GSK3β) and Rho-associated protein kinase (RhoK)) [16] at identified amino-acids residues. CRMP2 phosphorylation by (Cdk5), GSK3β, RhoK, or the Src-family kinases Fyn and Yes drives its diverse cellular functions, including neurite outgrowth, endocytosis, and ion-channel trafficking [16]. Using specific antibodies against CRMP2 and its phosphorylated forms, we quantified CRMP2 expression and phosphorylation levels in the mPFC after either extinction of cocaine self-administration or cue-induced reinstatement (after extinction). We compared results to respective Yoked saline controls and between tissue from extinction and cue-reinstated rats (**Fig. 3A,B**). Relative to Yoked saline controls, we found that CRMP2 expression level was decreased after extinction of cocaine self-administration, whereas subsequent exposure to cocaine-paired cues resulted in a potentiation of CRMP2 expression above extinction levels (One-way ANOVA, F_(27)_=14.06, p=0.0017, **Fig. 3C**). No changes were detected on CRMP2 phosphorylation levels on any of the tested sites (One-way ANOVAs for: p32: F_(27)_=0.0249; p509/p514: F_(27)_=0.9078; p522: F_(27)_=0.2148; p555: F_(27)_=0.3771**; Fig. 3D**). These results indicate that prior experience of cocaine self-administration and extinction decreased mPFC CRMP2 expression, while reinstatement potentiates CRMP2 expression levels.

**FIGURE 3:**
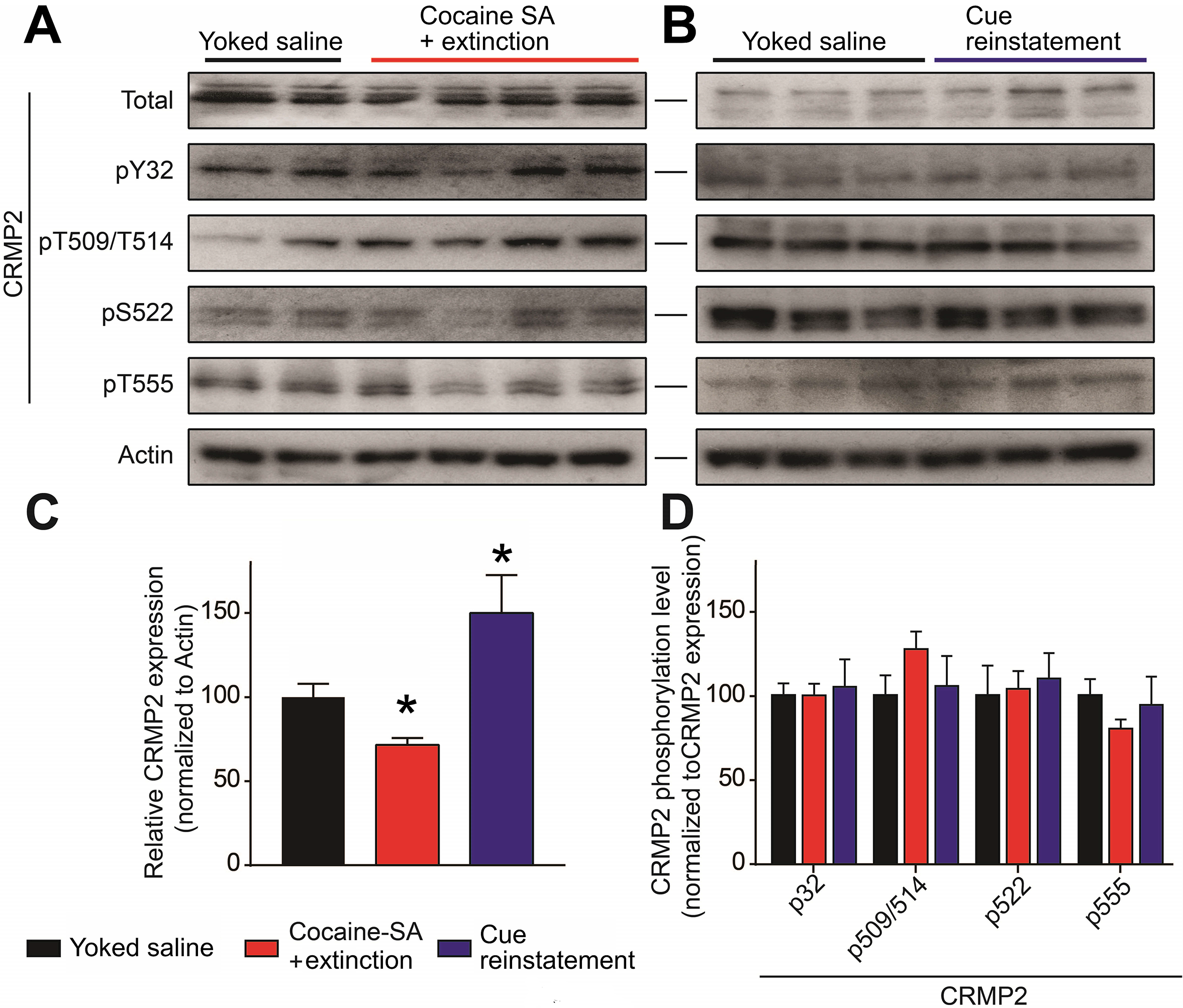
Cocaine SA alters the mPFC levels of CRMP2 protein expression, but not phosphorylation. (**A,B**) Representative Western blots showing treatment-related changes in mPFC CRMP2 expression and phosphorylation at Y32 (Fyn kinase), T509/T514 (GSK3β), S522 (Cdk5) and T555 (RhoK). Actin served as a loading control. Behavioral treatments (Yoked saline, Cocaine SA + extinction, Cue reinstatement) are described in Figure 2. (**C,D**) Summary bar graph showing differences in CRMP2 phosphorylation levels normalized as a fraction of the total CRMP2 expression level. (**C**) Relative to controls (Yoked saline), Cocaine SA + extinction treatment depressed CRMP2 protein expression (*p<0.05, Kruskall Wallis test). In rats with a history of Cocaine SA + extinction, Cue reinstatement potentiated CRMP2 expression relative to Cocaine SA + extinction (*p<0.05, Kruskall Wallis test.) (**D**) A comparison of phosphorylation across treatment groups showed no significant differences.

### CaV2.2 expression increased in the mPFC after cocaine reinstatement

CRMP2 is a regulator of the presynaptic voltage-gated calcium channel CaV2.2 [10; 34]. Because CRMP2 directly binds and regulates the expression of presynaptic CaV2.2 [8], decreased CRMP2 expression levels after extinction of cocaine self-administration would be expected to alter CaV2.2 expression. We used a specific CaV2.2 antibody to examine if CaV2.2 expression was altered in the mPFC of rats after extinction of cocaine self-administration or after cue-induced reinstatement (**Fig. 4A,B**). Compared to Yoked saline, we detected decreased CaV2.2 expression levels after extinction from cocaine self-administration (One-way ANOVA, F_(27)_=6.622, p=0.0446, **Fig. 4C**). In contrast, after cued reinstatement, CaV2.2 expression levels increased above control (Yoked saline) and extinction levels. Together with Fig. 3, these observations show concomitant differences in mPFC CRMP2 and CaV2.2 protein expression following both extinction of cocaine self-administration and cue-induced reinstatement. These results suggest that the CRMP2/CaV2.2 axis may contribute to cocaine reinstatement behaviors.

**FIGURE 4:**
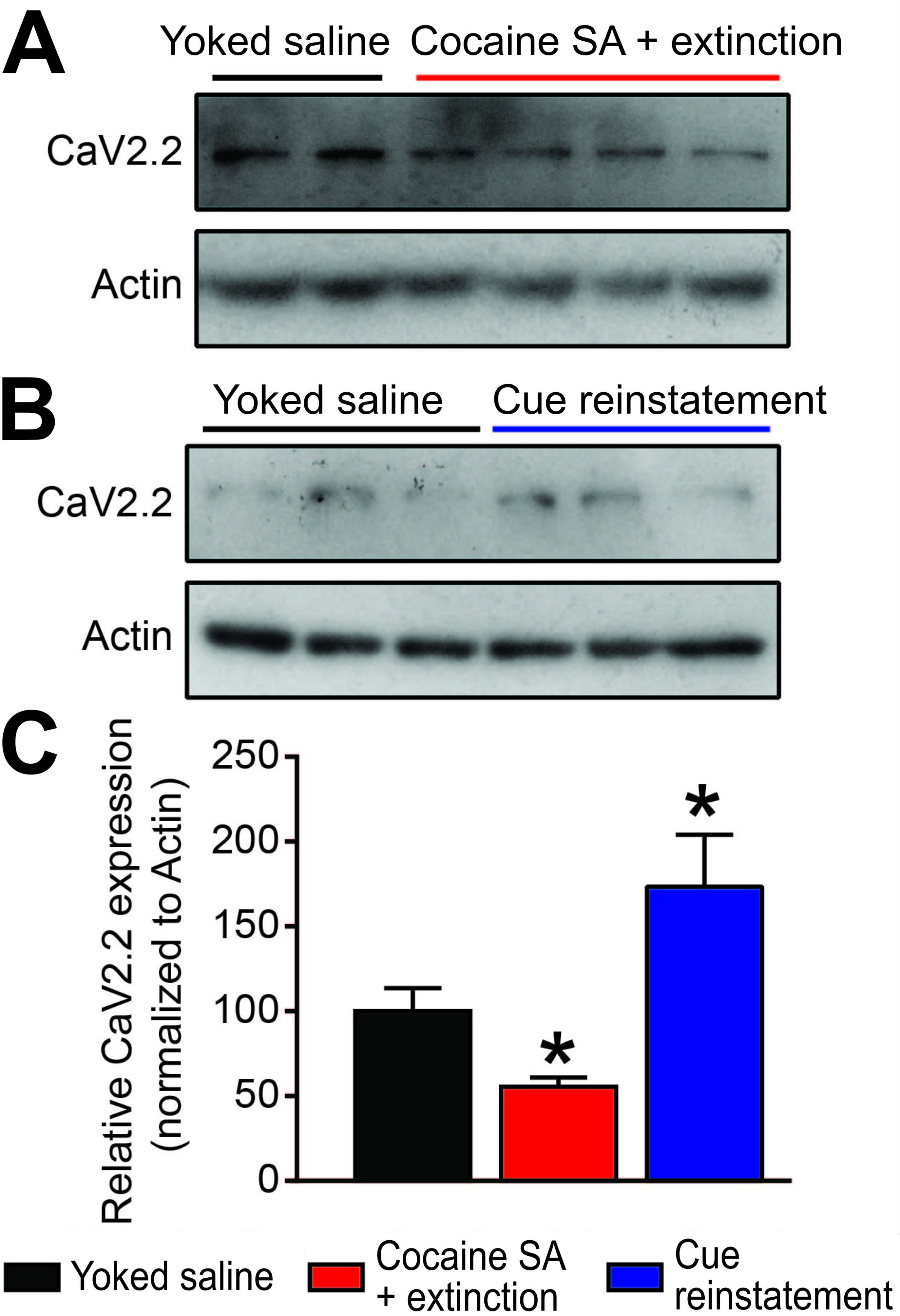
Cocaine SA + extinction and Cue reinstatement produce opposing actions on CaV2.2 expression. (**A,B**) Representative western blots showing treatment related changes in mPFC CaV2.2 expression. Actin served as a loading control. (**C**) Summary bar graph. Relative to Yoked saline controls, Cocaine SA + extinction decreased CaV2.2 expression (*p<0.05, Kruskall Wallis test). In contrast, relative to Cocaine SA + extinction, Cue reinstatement potentiated CaV2.2 expression (*p<0.05, Kruskall Wallis test).

### Inhibition of CRMP2 / Cav2.2 interaction decreases cue-induced reinstatement of cocaine seeking

To interrogate whether CaV2.2 regulation by CRMP2 could play a role in cocaine-seeking behavior, we utilized the CRMP2/CaV2.2 blocking peptide myr*-TAT-CBD3* [18]. This fifteen amino acid peptide has been shown to interfere with the interaction between CRMP2 and CaV2.2, resulting in disrupted surface colocalization of the two proteins and reduced calcium influx through CaV2.2 channels [18]. We previously reported that the peptide localizes to the plasma membrane of sensory neurons and is effective in blocking calcium influx within 5 minutes of application [44]. We first validated the ability of myr-TAT-CBD3 to inhibit the CRMP2/CaV2.2 interaction in brain lysates. We used a GST-pull down approach to detect CRMP2 interaction with CaV2.2 and βIII-tubulin as a positive control (**Fig. 5A**). Adding 30 μM of myr-TAT-CBD3 strongly inhibited CRMP2 interaction with CaV2.2 and increased CRMP2 interaction with βIII-tubulin (**Fig. 5B**), as we previously observed [44] (t-test, t_(6)_=9.895, p<0.0001).

**FIGURE 5:**
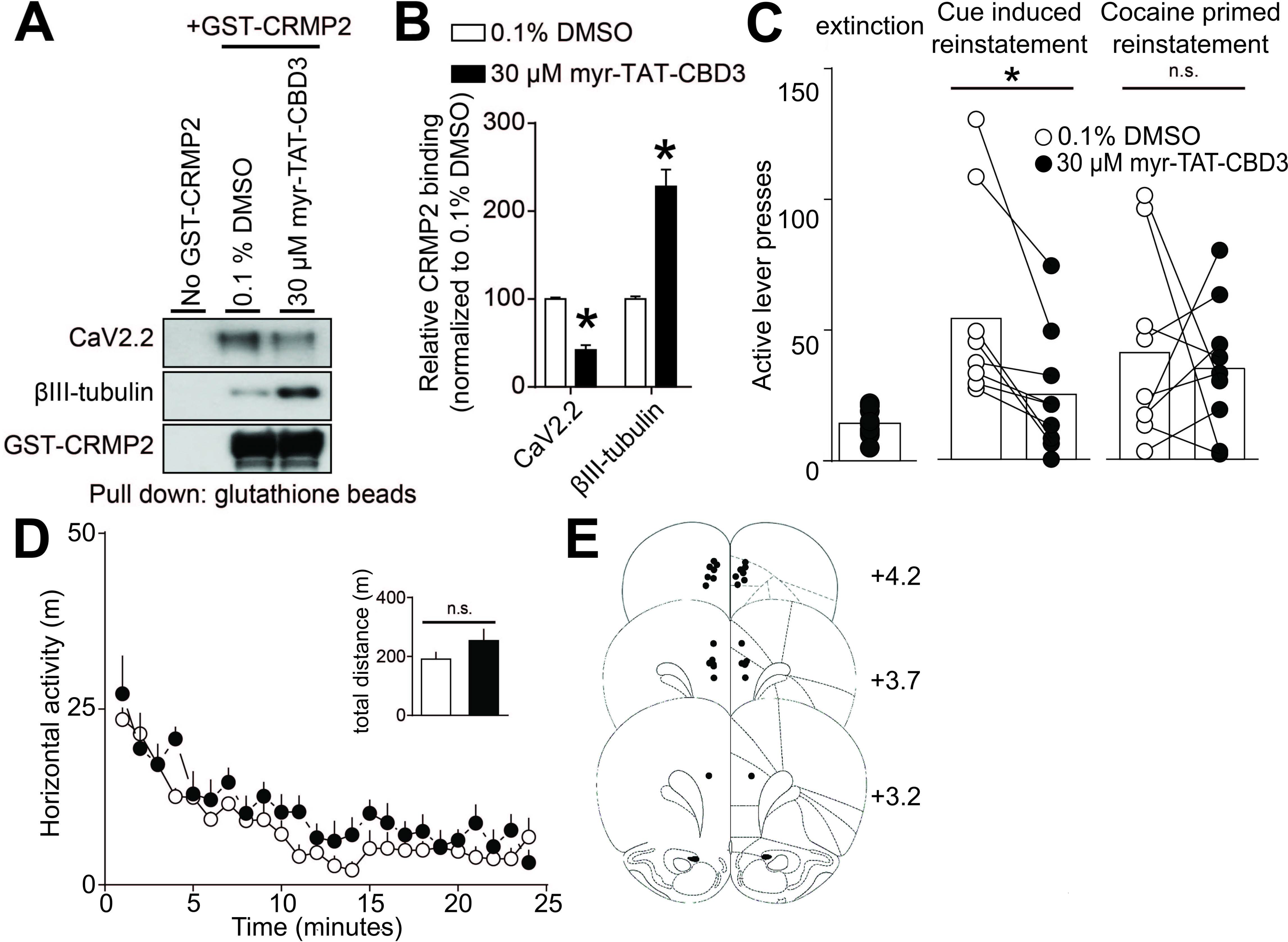
Inhibition of CRMP2-CaV2.2 interactions in the mPFC decreases reinstatement of cocaine seeking in response to drug-predictive cues, but not cocaine-prime. (**A**) A glutathione (GST)-pull down assay and Western analysis illustrating the interaction of CRMP2 with CaV2.2 and βIII-tubulin. Brain lysates were incubated with either myr-TAT-CBD3 or control vehicle. Minimal non-specific protein binding occurs in the no GST-CRMP2 condition (lane 1). (**B**) Bar graph summarizing the actions of vehicle or myr-TAT-CBD3 on the binding of CRMP2 to CaV2.2 or control βIII-tubulin (*p<0.05, Mann-Whitney test.). The myr-TAT-CBD3 peptide effectively inhibited CRMP2 interaction with CaV2.2 in brain lysate (n=7). (**C**) Bar graphs summarizing lever press responses during (*left*) extinction or different types of reinstatement testing (*middle*, cued-reinstatement; *right*, cocaine prime [10 mg/kg, i.p.]). All reinstatement testing (2 hr) occurred following bilateral mPFC infusion of vehicle or myr-TAT-CBD3. Relative to respective infusions of vehicle, myr-TAT-CBD3 decreased lever pressing in response to cocaine-associated cues (t_(8)_=3.665, p=0.0064), but not cocaine-prime (t_(8)_=0.3768, p=0.7161). (**D**) Relative to a bilateral mPFC infusion of vehicle, the infusion of myr-TAT-CBD3 does not reduce the spontaneous locomotor activity associated with a novel environment. The summary time course illustrates horizontal photocell counts (in 1 min bins) collected during the initial 25 min period of a 2-hr locomotor test session started 10 mins following mPFC drug infusion. *Inset* bar graph represents the total (mean ± SEM) horizontal activity for the 2-hr session (t_(8)_=1.343, p=0.2162). (**E**) Atlas plates summarize the location of microinjection cannula tips (Paxinos and Watson, 1998). Numbers denote distance from bregma in the anterior-posterior plane. For all infusions, vehicle was 0.1% DMSO and myr-TAT-CBD3 was applied at 30 μM.

We next used myr-TAT-CBD3 *in vivo* to examine if disrupting the interaction between CRMP2 and CaV2.2 reduced drug-seeking behavior. In cocaine self-administration-experienced rats extinguished of cocaine-seeking, we used a standard counterbalanced reinstatement design and microinjected myr-TAT-CBD3 or vehicle bilaterally into the mPFC 10 minutes prior to reinstatement testing. Each rat underwent 4 reinstatement sessions separated by at least two days of extinction, counterbalanced with respect to infusion of either myr-TAT-CBD3 or vehicle. Compared to vehicle infusion, mPFC infusion of myr-TAT-CBD3 significantly reduced active-lever pressing during cue-induced reinstatement testing (t_(8)_=3.665, p=0.0064, **Fig. 5C**). Active lever presses induced by a priming cocaine injection (10 mg/kg, i.p.) did not differ between myr-TAT-CBD3 and vehicle reinstatement tests (t_(8)_=0.3768, p=0.7161, **Fig. 5C**). Subsequent bilateral mPFC infusion of myr-TAT-CBD3 did not reduce spontaneous locomotor activity in a novel chamber (t_(8)_=1.343, p=0.2162), suggesting the myr-TAT-CBD3 action on cue-induced reinstatement was not due to a nonspecific reduction in motor behavior. Histological confirmation of mPFC cannula placements are shown in Fig 5E.

## Discussion

Here, we show that a history of prior extinction from cocaine self-administration alters the subsequent regulation of CaV2.2 in the mPFC and its binding partner the collapsin response mediator protein-2 (CRMP-2), a key protein regulating presynaptic calcium channels and neurotransmitter release. Cocaine (but not saline) self-administration and extinction training reduced expression of both CaV2.2 and CRMP2. Other studies, not using extinction, also found decreased CRMP2 expression 24-hr after a final contingent of cocaine SA [50] or after daily noncontingent methamphetamine administration [25]. Therefore, the reductions we found appear persistent despite extinction training. Re-exposure to cues predictive of cocaine robustly enhanced both CRMP2 and CaV2.2 expression relative to extinction levels. Furthermore, disruption of the CRMP2-CaV2.2 protein-protein interaction in the mPFC decreased cue-induced (but not cocaine-primed) reinstatement of cocaine seeking. The cellular mechanism underlying these changes in expression remain unknown. The breadth of CRMP2 protein interactions are numerous and widespread, yet our results suggest that in the PFC, CRMP2 interactions with CaV2.2 are critical for cued-reinstatement of cocaine, an animal model of relapse.

To the best of our knowledge, no existing studies have directly examined CRMP2 function in PFC circuitry in the context of drug-seeking. Recent studies indicate that abused drugs or their associated behaviors dysregulate CRMP2, glutamate release, and VGCCs. In rats with a history of Cocaine SA, re-exposure to cocaine-paired cues enhances glutamate release in the mPFC [46]. A similar response occurs in the PFC with non-contingent methamphetamine exposure, but not in animals with a history of sucrose SA [31]. Thus, some have suggested that drugs of abuse (but not natural rewards) can sensitize the responsiveness of glutamate release in the mPFC [46; 47].

Adaptations in presynaptic glutamate release could reflect altered expression of VGCCs mediated through changes in the α2δ-1 subunit that regulates recycling of CaV2.2 to pre-synaptic sites [12; 24; 51]. Repeated noncontingent methamphetamine [21] or morphine [45], as well as contingent cocaine administration [48] increases α_2_δ expression in the PFC and nucleus accumbens. Conversely, the clinical therapeutic Gabapentin, which binds α_2_δ and decreases CaV2.2 channel currents and neurotransmitter release [23], reverses some of these actions. Gabapentin decreases the probability of transmitter release in the accumbens in cocaine-extinguished rats but not control (Yoked saline) animals [48]. Intracerebroventricular delivery of gabapentin reduces the sensitization and conditioned place preference associated with both methamphetamine [28] and morphine [45]. Such findings highlight the potential efficacy of direct targeting of the CaV2.2 system for various substance abuse disorders. However, targeting VGCC or α2δ-1 has the potential for significant side effects, such as disruption of synapse formation [17]. There remains a need for better alternatives.

### Indirectly targeting CaV2.2: CRMP2 phosphorylation

Targeting the regulation of CaV2.2 channels may potentially lessen adverse effects associated with direct channel block. An alternative strategy to ‘fine tune’ calcium channel density at the glutamate terminal is to target CRMP2. Either reducing the effects of Cocaine SA on the phosphorylation of CRMP2 or inactivating CRMP2 may be beneficial. The first approach is challenging, as CRMP2 is a multi-phosphorylated protein in neurons [2; 53]. The second approach for inactivation may be more feasible, in that CaV2.2 binds to the non-phosphorylated but active form of CRMP2 [11] that can be inactivated with the myr-TAT-CBD3 peptide.

On this basis, we examined phosphorylation sites known to regulate CRMP2 function and selected 5 (other than the Yes site) that are known to be sensitive to cocaine: Yes, Fyn, GSK3β, Cdk5 and Rho [3; 42]. We predicted that phosphorylation of CRMP2 would be enhanced after Cocaine SA and extinction or reinstatement. However, we found no differences between treated and control animals. Future studies will be required to evaluate whether extinction training could explain this lack of effect. Considering that all reports mentioned above share a common feature of animals with recent or immediate access to cocaine, it is possible that cocaine-related changes in kinase activity are transient and resolve over time. Alternatively, a different kinase, not included in our study, may be involved (e.g., PKA), as reported in the nucleus accumbens [6]. A kinase-based strategy may be helpful to treat ongoing drug self-administration, but based on our present findings, may be less helpful for prevention of relapse after the cessation of cocaine self-administration.

### Indirectly targeting CaV2.2: peptidomimetic inactivation of CRMP2

Our current data indicated that both cocaine self-administration and reinstatement robustly altered the expression of CRMP2 and CaV2.2 in the mPFC. The direction of the response (decreases or increases, respectively) depended on the behavioral condition. Imaging studies in humans report widespread PFC hypoactivity following cessation of cocaine [19; 20]. It is tempting to speculate that this PFC hypoactivity may coincide with our observations of reduced CaV2.2 expression following extinction from Cocaine SA. To the best of our knowledge, however, there are no direct supporting neurochemical studies in rodents correlating prior cocaine exposure with reductions in basal transmitter release. Electrophysiology studies examining synaptic transmission have reported both increases [37] and decreases [39] in EPSC frequency in the prelimbic mPFC following cocaine place preference training. Meanwhile, others have not observed changes in basal excitatory synaptic strength in the mPFC following non-contingent cocaine exposure, despite changes in post-synaptic plasticity [41]. Future studies are required to determine whether the observed changes in CRMP2/CaV2.2 proteins contribute to adaptations in presynaptic neurotransmitter release following extinction from Cocaine SA.

Reinstatement of drug seeking, in contrast, has been associated with increased activity in the PFC [9; 19]. Because we observed increased CaV2.2 expression following cue-induced reinstatement, we tested if a functional CRMP2-CaV2.2 protein-protein interaction in the PFC was required for cued-reinstatement. We targeted CaV2.2 indirectly with a peptide derived from the CRMP2 backbone, myr-TAT-CBD3, shown previously not to impact memory retrieval, motor coordination, depression-associated behaviors [7] or CPP [18] after intracerebroventricular administration. We also independently confirmed that direct PFC infusion of myr-TAT-CBD3 produced no measurable change in nonspecific movements. When applied locally to the PFC, this peptide reduced both CRMP2 binding to CaV2.2 and reinstatement in response to drug-cues, but not to cocaine-priming.

A recent study demonstrated that knockdown of CRMP2 in the nucleus accumbens or systemic administration of a CRMP2 antagonist, (*R*)-lacosamide (Vimpat), suppresses ongoing excessive drug (alcohol) intake [29]. This (*R*)-enantiomer of lacosamide effectively blocks CRMP2-dependent microtubule formation, but is without action on CaV2.2 channel function or CRMP2 phosphorylation states, the regulatory event driving CRMP2-CaV2.2 association [32]. Combined with the data from our study, one scenario that emerges is that CRMP2 regulation of microtubule formation contributes to the initial or ongoing stages of excessive drug taking. Following the cessation of repeated self-administration, transient phosphorylation of CRMP2 may then drive an increased association with CaV2.2, resulting in trafficking of this complex to terminals to support long-lasting structural plasticity. Therefore, targeting the CaV2.2-CRMP2 protein-protein interaction may be most effective at eliminating relapse behavior once drug-seeking behavior is already learned.

In summary, our work highlights the instrumental role of CRMP2 coupling to CaV2.2 in cocaine reinstatement behaviors. (*S*)-Lacosamide, edonerpic maleate [1], and other promising candidate CRMP2 targeting compounds warrant further investigation and characterization for therapeutic potential.

## Acknowledgements

NIDA grants F31-DA03689 (W.C.B.), T32-DA007288 (W.C.B.; PD: J.F.M.), R01-DA033342A (A.C.R.), P50-DA015369 (PI: A.C.R.; PD: P.W.K.), R01 DA046476 (A.C.R), R01-DA042852 (R.K.), and NINDS grant R01-NS098772 (R.K.).

## Author Contributions

W.C.B., A.M., B.H., R.K., and A.C.R designed research. W.C.B., A.M., B.H., C.G.K., A.S. performed research and analyzed data. W.C.B., A.M., R.K. and A.C.R wrote the paper.

